# Redundancy in the central tachykinin systems safeguards puberty onset and fertility

**DOI:** 10.1101/443333

**Authors:** Silvia León, Chrysanthi Fergani, Rajae Talbi, Serap Simavli, Caroline A. Maguire, Achi Gerutshang, Stephanie B. Seminara, Víctor M. Navarro

## Abstract

The tachykinin neurokinin B (NKB, *Tac2*) is critical for GnRH release. NKB signaling deficiency leads to infertility in humans. However, some patients reverse this hypogonadism resembling the fertile phenotype of *Tac2*KO and *Tacr3*KO (encoding NKB receptor, NK3R) mice despite the absence of NKB signaling. Here, we demonstrate that in the absence of NKB signaling, other tachykinins (substance P and neurokinin A [NKA], encoded by *Tac1*) may take over to preserve fertility. The complete absence of tachykinins in *Tac1/Tac2*KO mice leads to delayed puberty onset in both sexes and infertility in 80% of females (but not males), in contrast to the 100% fertile phenotype of *Tac1*KO and *Tac2*KO mice separately. Furthermore, we demonstrate that NKA controls puberty onset and LH release through NKB-independent mechanisms in the presence of sex steroids and NKB-dependent mechanisms in their absence. In summary, tachykinins interact in a coordinated manner to ensure reproductive success in female mice.

## Introduction

Tachykinins (TACs) are a large family of peptides that includes neurokinin A (NKA) and substance P (SP), encoded by *TAC1*, and neurokinin B (NKB), encoded by *TAC3* (or *Tac2* in rodents) (1). TACs act on different G protein-coupled receptors: NK1R (encoded by *Tacr1*), the receptor for SP; NK2R (*Tacr2*), the receptor for NKA; and NK3R (*Tacr3*), the receptor of NKB. These TAC systems are expressed throughout the central nervous system, where they participate in a variety of physiological functions, e.g. nociception and fear conditioning (1, 2).

Recently, the NKB/NK3R system has emerged as a critical neuroendocrine regulator of reproductive function. A growing body of evidence from our lab and others has documented the stimulatory role of NKB on GnRH release in an estradiol and kisspeptin dependent manner in all studied species, including the human (3).

Moreover, inactivating mutations in *TAC3/TACR3* genes lead to hypogonadotropic hypogonadism (HH) in patients, presenting delayed or absent puberty onset and infertility (4, 5). However, a number of human patients bearing these mutations overcome initial pubertal failure and central hypogonadism, with a later ‘awakening’ of GnRH secretion and hypogonadism reversal (6); a phenotype that resembles that of *Tac2*KO and *Tacr3*KO mice, which are sub-fertile (7, 8). The underlying mechanism responsible for the reversal phenotype of these patients remains unknown. In addition to NKB, the SP/NK1R system also participates in the central regulation of the gonadotropic axis, likely via kisspeptin-dependent mechanisms (9, 10). Supporting this contention, (*a*) studies in humans, rabbits and rats showed a central stimulatory role of SP on LH release (11-13); (*b*) Kiss1 neurons are activated by SP (9); (*c*) SP mRNA and protein has been found in the ARC of rodents (10, 14); (*d*) SP immunoreactivity has been detected in Kiss1 and NKB neurons in the human infundibular nucleus (15); (*e*) chronic SP administration advances puberty onset in rodents (16) and (*f*) *Tac1*KO mice with congenital absence of SP, display delayed puberty onset and reproductive impairments (16, 17).

Interestingly, because *Tac1* encodes both SP and NKA we cannot rule out that the reproductive phenotype observed in *Tac1*KO mice (delayed puberty onset and sub-fertility) (16, 17) is not, at least in part, due to the absence of NKA signaling, whose participation in the control of puberty onset and fertility remains largely unknown. We and others have documented that NKA also induces LH release in rodents (10, 18, 19) in a kisspeptin-dependent manner (10). Interestingly, the stimulatory action of NKA on LH release is dependent on the presence of physiological levels of circulating sex steroids, while in their absence, NKA inhibits LH release, similar to NKB (10). However, unlike NKB, NKA’s receptor (NK2R) is not present on Kiss1 or GnRH neurons (10). We therefore hypothesize that NKA must act upstream of Kiss1 neurons, on an unknown population of neurons, that in turn control NKB release. Alternatively, all TAC ligand-receptor systems have been reported to display cross-reactivity (20), implicating that cross-activation of NK3R by NKA could be an explanation for the NKB-like action of NKA. Overall, there is compelling evidence that all TACs (not only NKB) participate in the control of GnRH release. This highlights the importance of deciphering the mechanism of action for each one individually, as well as in combination with other tachykinins, in order to better understand their role in the neuroendocrine control of reproduction.

## Results

### 1. Complete absence of tachykinins in double *Tac1/Tac2* KO females severely disrupts puberty onset and fertility in females

In order to test our hypothesis, that there is redundancy in the tachykinin systems that preserves fertility in the absence of NKA/SP signaling (*Tac1*KO mice) or NKB signaling (*Tac2*KO and Tacr3KO mice) individually (7, 8, 16), we generated double *Tac1/Tac2* KO mice and compared their reproductive maturation and fertility to their single mutant littermates. Interestingly, while *Tac1*KO and *Tac2*KO mice displayed delayed VO, as reported previously, double *Tac1/Tac2* KO females presented only a moderate (not significant) delay in VO compared to WT littermates (**Figure 1 A-C**), however, they failed to show any signs of first estrous for over 30 days post VO (**Figure 1 D**). Of note, first estrous is a more accurate marker of puberty onset as it indicates central activation of the gonadotropic axis. Furthermore, in a fertility test in which adult females were mated with proven fertile WT males, only 20% of *Tac1/Tac2* KO females were able to deliver pups over a 90-day long period of mating (**Figure 1 E**). Moreover, the parturition latency of these small portion of *Tac1/Tac2* KO females was longer than in WT controls (WT: 21.66 ± 0.66 days; *Tac1/Tac2* KO: 29.5 ± 0.5 days; **p<0.01) (**Figure 1 F**) and the litter size was significantly smaller than in controls (WT: 7.33 ± 1.08 pups; *Tac1/Tac2* KO: 4 ± 0 pups; **p<0.01) (**Figure 1 G**). This largely infertile phenotype of *Tac1/Tac2* KO females is also supported by the reduced number of corpora lutea in their ovaries (WT: 3.5 ± 0.28 CL/ovary; *Tac1/Tac2* KO: 0.75 ± 0.25 CL/ovary; ***p<0.001) (**Figure 1 H, I**) and significantly lower ovarian weight than controls (WT: 18.50 ± 1.70 mg; *Tac1/Tac2* KO: 10.83 ± 1.63 mg; **p<0.01). Moreover, since previous studies had documented an improvement of estrous cyclicity with age in *Tac2*KO mice (8), we assessed whether this was also the case in *Tac1/Tac2* KO females at a young (3 months) and older (8 months) age and observed that, in both cases, KO females failed to show any signs of regular estrous cyclicity (**Figure 1 J,K**).

**1.**
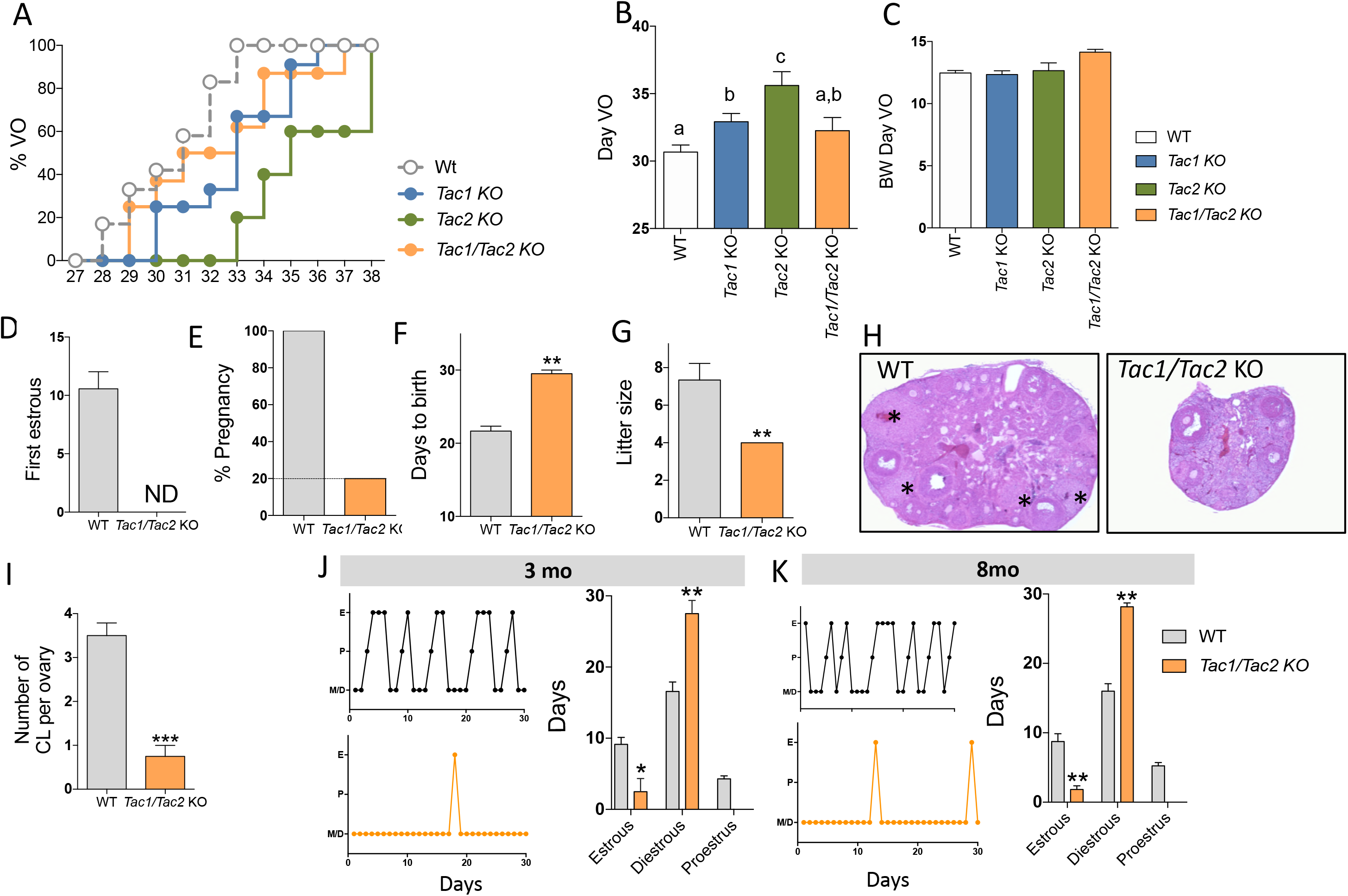
Complete absence of tachykinins in double *Tac1/Tac2* KO females severely disrupts puberty onset and fertility in females. **(A)** Accumulated percentage of mice with vaginal opening (VO), an indirect assessment of puberty onset by daily visualization for two weeks post weaning (WT n=12; *Tac1* KO n=12;*Tac2 KO* n=5 and *Tac1/Tac2* KO n=8). **(B)** Mean postnatal day of VO and **(C)** BW the day of VO (WT n=12; *Tac1* KO n=12; *Tac2* KO n=5 and *Tac1/Tac2* KO n=8). Different letters indicate statistically different values (2 Way ANOVA followed by Newman Kleus post-hoc test, p<0.05). **(D)** Mean postnatal day of first estrous as determined by histology samples taken over the course of 30 days after VO (WT n=9 and *Tac1/Tac2* KO n=8). **(E)** Fertility test in which adult WT and *Tac1/Tac2* KO females were mated with proven fertile WT males (WT n=3 and *Tac1/Tac2* KO n=10). Percentage of females that delivered pups. **(F)** Number of pups per litter for one set of breeding pairs over 90 days of ongoing breeding is significantly lower in *Tac1/Tac2* KO. **p<0.01 Student t-test. **(G)** Parturition latency in WT (n = 3) and *Tac1/Tac2* KO (n = 10) mice. **p<0.01 Student t-test. **(H)** Ovarian histology of WT (n = 3) and *Tac1/Tac2* KO mice (n = 10). Each asterisk denotes a corpus luteum. **(I)** Number of corpora lutea in the middle section of the ovary from WT (n=4) and *Tac1/Tac2* KO (n=4) female mice. ***p<0.001 Student t-test. **(J)** Representative examples of daily estrous cycle phases over 30 days in 3-month-old WT (n=7) and *Tac1/Tac2* KO (n=8) female mice. Days on estrous phase (E), proestrous phase (P) and diestrous phase (D). WT and *Tac1/Tac2* KO. *p<0.05 Student t-test; **p<0.01 Student t-test. **(K)** Representative examples of daily estrous cycle phases over 30 days in 8-month-old WT (n=4) and *Tac1/Tac2* KO (n=6) female mice. Days on estrous phase (E), proestrous phase (P) and diestrous phase (D). WT and *Tac1/Tac2* KO. *p<0.05 Student t-test; **p<0.01 Student t-test.

### 2. *Tac1/Tac2* KO females display disrupted LH pulses and response to ovariectomy

To further characterize the reproductive phenotype of *Tac1/Tac2* KO females, LH pulsatility and long-term response to OVX were assessed. As expected, OVX *Tac1/Tac2* KO females displayed a severely disrupted LH pulse pattern (fewer pulses) over 150 minutes compared to OVX WT (WT: 3.33 ± 0.56 pulses/150min; *Tac1/Tac2* KO: 1.86 ± 0.26 pulses/150min; *p<0.05) (**Figure 2 A-C**). Interestingly, when the response of LH to OVX was assessed, we observed a biphasic response in which LH levels were reduced compared to WT in *Tac1/Tac2* KO and *Tac2*KO females 2 days post OVX; however, there was a quick rebound that led to normal LH levels in both KO models compared to WT females at 7 days post OVX (**Figure 2 D**). Next, we aimed to assess the ability of *Tac1*KO, *Tac2*KO and *Tac1/Tac2* KO females to respond to the central administration of kisspeptin (Kp-10) and GnRH. As we previously reported, the lack of *Tac1* led to a diminished LH response to Kp-10 (17), which was replicated in *Tac1/Tac2* KO females and extended to the GnRH response (**Figure 2 E, F**), while the expression of the tachykinin receptors (*Tacr1*, *Tacr2* and *Tacr3*), *Kiss1* and *Pdyn* remained unaffected (**Figure 2 G, H**).

**2.**
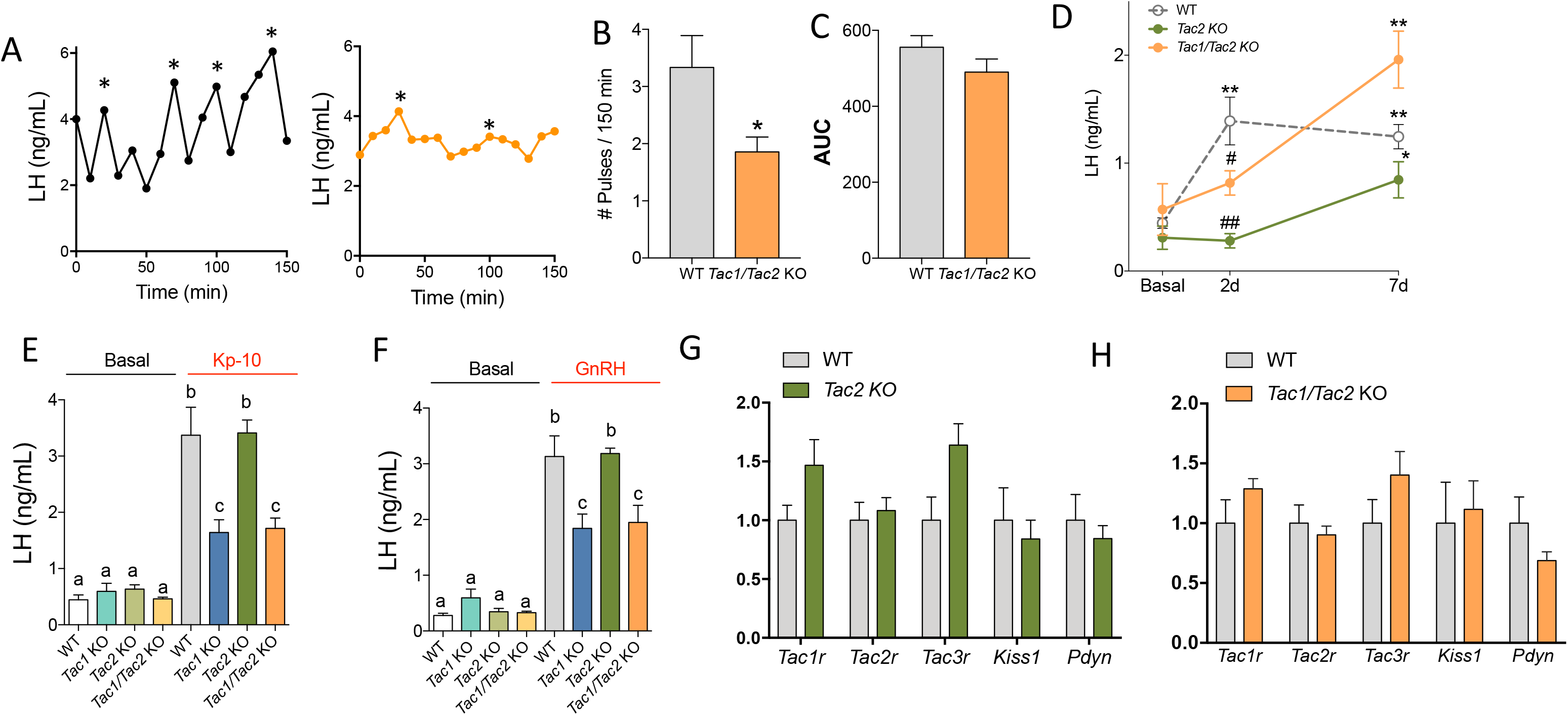
*Tac1/Tac2* KO females display disrupted LH pulses and increased response to ovariectomy over 21 days. **(A)** Representative pattern of LH pulsatility over 150 min (WT n=6 and *Tac1/Tac2* KO n=7). Analyzed using a MATLAB-bases algorithm. **(B)** Number of pulses over a 90 minutes sampling period. **p<0.01 Student t-test. **(C)** Area under the curve (AUC) of the pattern of LH pulsatility over 150 min. Student t-test. **(D)** Circulating LH levels before and 2d, 7d, 14d and 21d (days) after bilateral ovariectomy (OVX). WT (n=9), *Tac2* KO (n=4) and *Tac1/Tac2* KO (n=6) *p < 0.05; **p < 0.01 vs. corresponding basal (pre-OVX) levels; 2 Way ANOVA followed by Tukey post hoc test; #p < 0.05; ##p < 0.01, KO vs. corresponding WT at each specific time point. **(E)** Circulating LH levels 15 min after the injection of kisspeptin-10 (50 pmol/5ul/icv). WT (n=7), Tac1 KO (n=8), Tac2 KO (n=5) and *Tac1/Tac2* KO (n=7). Different letters indicate statistically different values (2 Way ANOVA followed by Tukey post-hoc test, p<0.05). **(F)** Circulating LH levels 30 min after the injection of GnRH (0.25 ug/100μl/ip). WT (n=7), *Tac1* KO (n=8), *Tac2* KO (n=5) and *Tac1/Tac2* KO (n=7). Different letters indicate statistically different values (2 Way ANOVA followed by Tukey post hoc test, p<0.05). **(G, H)** Expression of *Tac1r, Tacr2, Tacr3, Kiss1* and *Pdyn* in the mediobasal hypothalamus of *Tac2*KO **(G)** and *Tac1/Tac2*KO **(H)** females and their WT controls. WT (n=7), *Tac2* KO (n=9) and *Tac1/Tac2* KO (n=7).

### 3. Complete absence of tachykinins in double *Tac1/Tac2* KO males delays puberty onset but does not affect fertility

Similar to the delay in first estrus in female *Tac1/Tac2* KO mice, males displayed delayed puberty onset as shown by the timing of preputial separation (**Figure 3 A-C**). However, and unlike females, *Tac1/Tac2* KO males were fertile, with a 100% success ratio in their ability to impregnate WT females (**Figure 3 D-F**). Concordantly, their testicular histology was normal compared to controls, showing mature sperm in all cases (**Figure 3 G**) and normal testicular weight (WT 163.31 ± 6.38 mg; *Tac1/Tac2* KO 172.84 ± 9.20 mg). As in females, *Tac1/Tac2* KO males presented a delay in the response to gonadectomy 2 days after surgery, however, LH levels returned to normal by 7d post GDX (**Figure 3H**). Finally, similar to our previous reports in *Tac1*KO males (17) and to our present data in females (**Figure 2**), *Tac1/Tac2* KO males have a significant decrease in the response to Kp-10 and GnRH (**Figure 3 I, J**).

**3.**
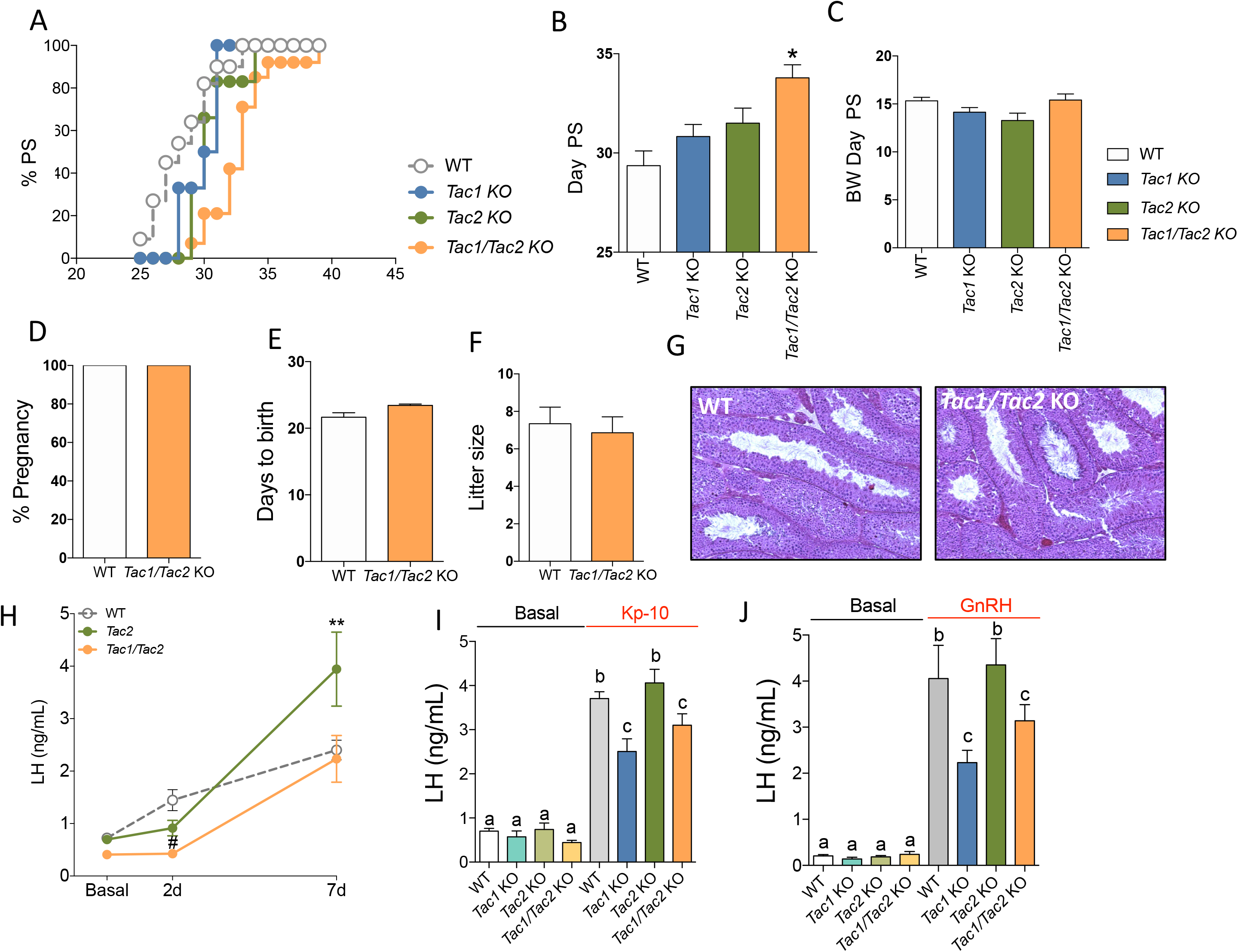
Complete absence of tachykinins in double *Tac1/Tac2* KO males delays puberty onset without affecting fertility. **(A)** Accumulated percentage of mice with preputial separation (PS) as an indirect marker of puberty onset for two weeks post weaning, WT (n=11), *Tac1* KO (n=6), *Tac2* KO (n=6) and *Tac1/Tac2* KO (n=14). **(B)** Mean postnatal day of PS and **(C)** BW the day of PS (WT n=12; *Tac1* KO n=12; *Tac2* KO n=5 and *Tac1/Tac2* KO n=8). **(D)** Percentage of success to impregnate WT females (WT n=3 and *Tac1/Tac2* KO n=7). **(E)** Number of pups per litter over 90 days of breeding and **(F)** parturition latency. **(G)** Testicular histology was normal in *Tac1/Tac2* KO compared to controls, showing mature sperm in all cases (WT n=4 and *Tac1/Tac2* KO n=4). **(H)** Circulating LH levels before and 2d, 7d, 14d and 21d (days) after bilateral gonadectomy (GNX). WT (n=13), *Tac2* KO (n=4) and *Tac1/Tac2* KO (n=10); **p < 0.01 vs. corresponding basal (pre-GNX) levels; #p < 0.05, KO vs. corresponding WT at each specific time point. **(I)** Circulating LH levels 15 min after the injection of kisspeptin-10 (50 pmol/5ul/icv). WT (n=20), *Tac1* KO (n=7), *Tac2* KO (n=12) and *Tac1/Tac2* KO (n=18). Different letters indicate statistically different values (2 Way ANOVA followed by Tukey post hoc test, p<0.05). **(J)** Circulating LH levels 30 min after the injection of GnRH (0.25 ug/100μl/ip). WT (n=7), *Tac1* KO (n=4), *Tac2* KO (n=8) and *Tac1/Tac2* KO (n=8). Different letters indicate statistically different values (2 Way ANOVA followed by Tukey post hoc test, p<0.05).

### 4. The receptor of NKA (NK2R) is expressed in VMH *Tac1* neurons

Our previous studies have documented the existence of mRNA of the NKB receptor (NK3R) and SP receptor (NK1R) in ~100% and ~50%, respectively, of Kiss1 neurons in the ARC, while the receptor of NKA (NK2R) was undetectable in Kiss1 or GnRH neurons (10). We therefore aimed to assess if NK2R (encoded by *Tacr2*) is expressed in other ARC neurons or in its nearby VMH nucleus and whether it colocalizes with Tac1 neurons in these areas, as a critical source of SP and NKA in the control of kisspeptin/GnRH release. Our *in situ* hybridization (RNAscope) results showed that *Tacr2* is expressed in both ARC and VMH nuclei and colocalizes with Tac1 neurons in the VMH of adult WT mice regardless of the E2 milieu (**Figure 4**).

**4.**
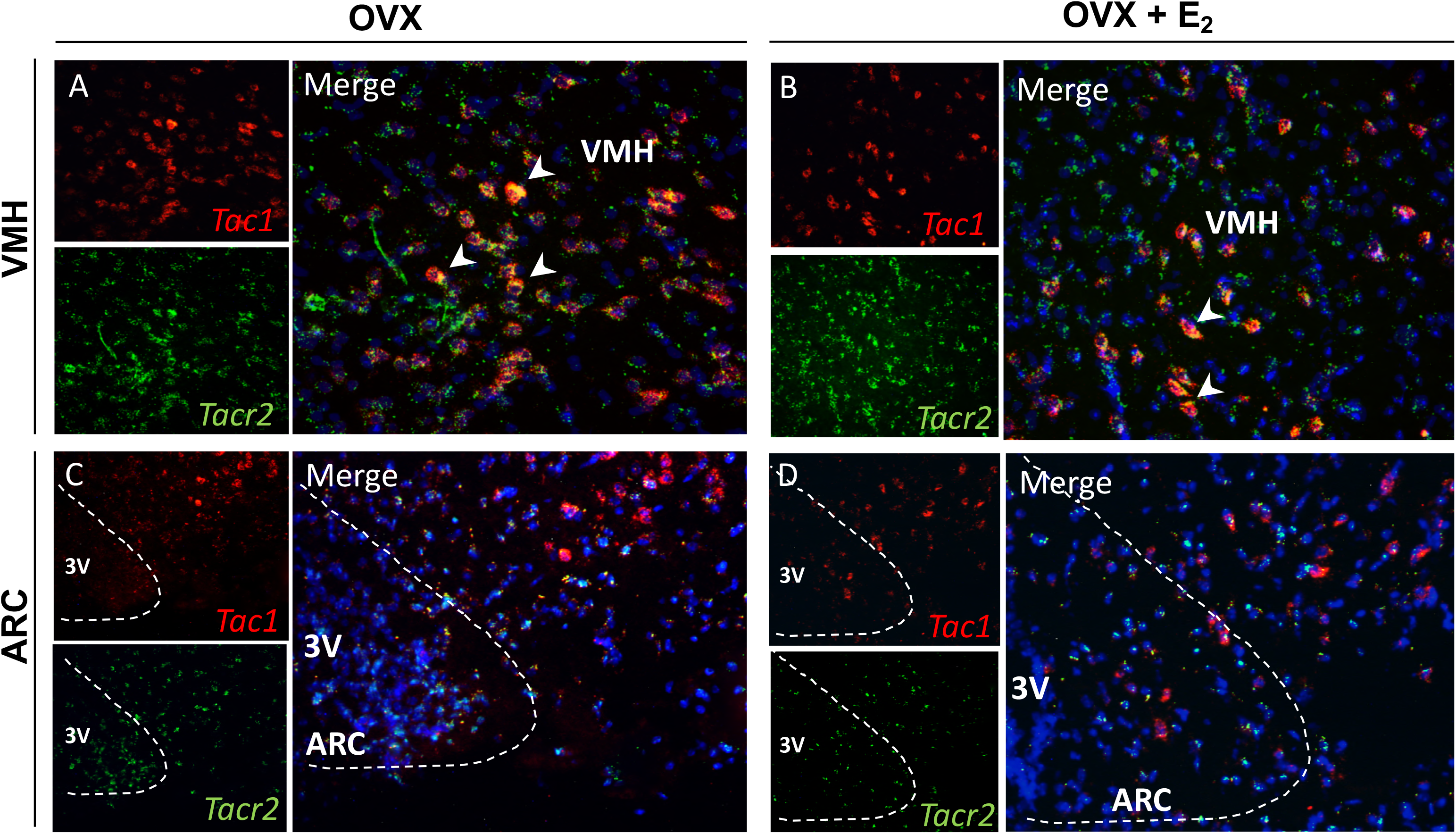
The receptor of NKA (NK2R) is expressed in VMH Tac1 neurons. Representative double label ISH depicting co-localization of *Tac1* (red) and *Tacr2* (green) mRNA. **(A)** VMH and **(C)** ARC of female WT C57Bl/6 mice after 1 week of OVX. **(B)** VMH and **(D)** ARC of female WT C57Bl/6 OVX mice after 1 week of E_2_ replacement.

### 5. Advancement of puberty onset after chronic activation of NK2R in female mice

Figures 1 and 3 evidenced that tachykinins are needed for normal timing of puberty onset in male and female mice. Likewise, our previous studies have demonstrated that chronic administration of specific agonists of the NK1R and NK3R receptor are able to advance puberty onset in rodents (16, 21), indicating that these systems are in place before puberty and likely participate in the proper timing of puberty onset. However, whether NKA/NK2R is also involved in the re-awakening of the gonadotropic axis at the time of puberty is unknown. To address this question, we chronically (every 12h) treated WT females with a specific agonist of NK2R from weaning age (22d) to 32d. We observed that this treatment was able to advance puberty onset as evidenced by the advanced timing of VO and increased uterus and ovarian weight compared to controls (Uterus weight: Control: 19.55 ± 1.89 mg; NK2R-Ag treated: 23.79 ± 1.61 mg; *p < 0.05. Ovarian weight: Control: 8.3 ± 0.56 mg; NK2R-Ag treated: 11.7 ± 0.44 mg; ***p < 0.001) (**Figure 5 A-F**).

**5.**
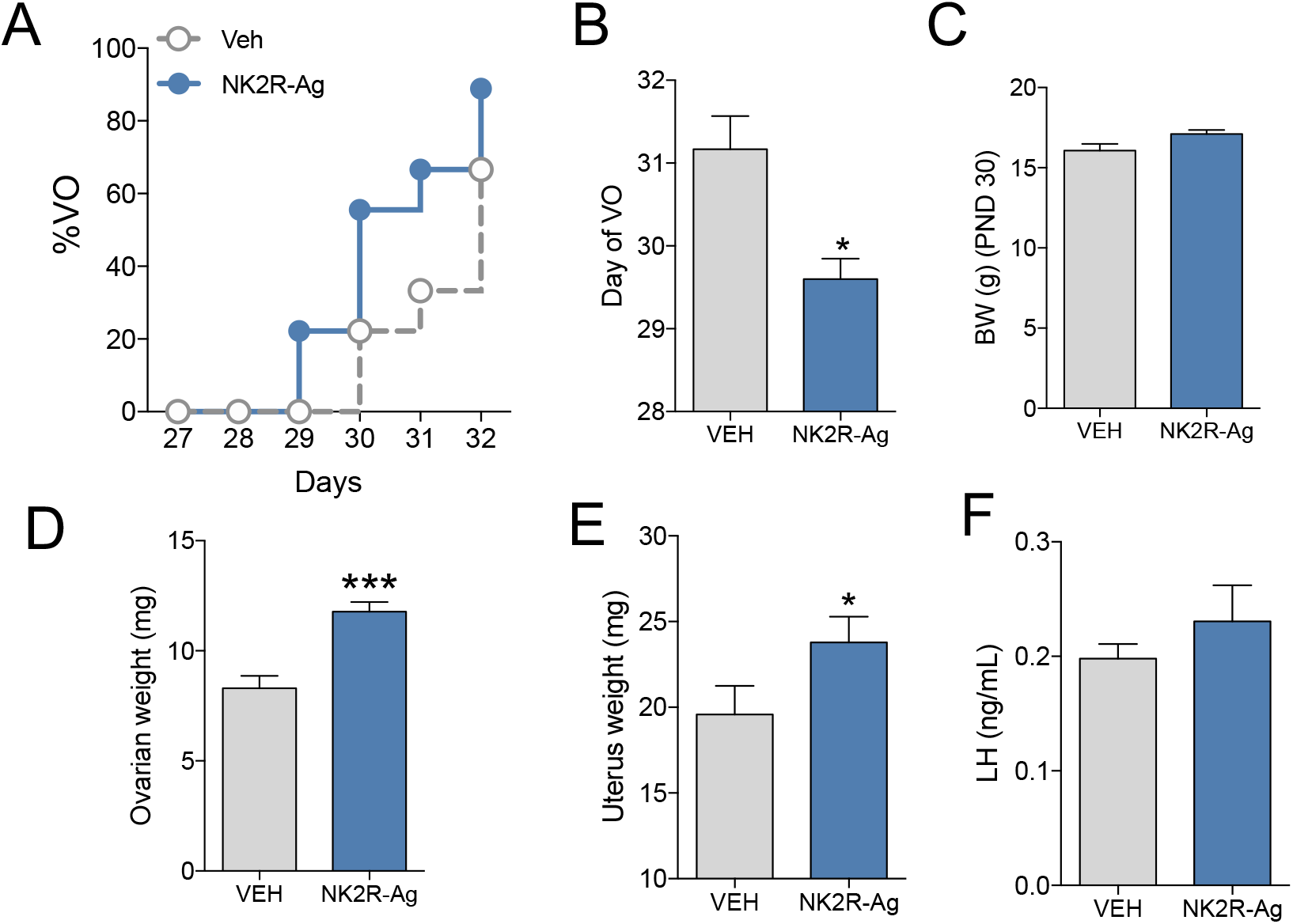
Advancement of puberty onset after chronic activation of NK2R in female mice. Repeated stimulation (every 12 h) of WT female mice with GR64349 (NK2R-A, 3 nmol/100ul/ip) or vehicle (0.9% NaCl/100ul/ip) from p23 to p32 (n≥6 per group). **(A)** Progression of VO, **(B)** mean postnatal day of VO and **(C)** BW the day of 50% of the control animals displayed VO. **(D)** Uterine weight, **(E)** ovarian weight and **(F)** serum LH levels at p32. Statistical analysis was performed using a 2-tailed t test (*p < 0.05; ***p< .001).

### 6. The stimulatory action of NKA is independent of NKB but dependent of kisspeptin

We previously reported that NKA and NKB action on LH release is largely equivalent, i.e. both increase LH release in the presence of physiological circulating levels of E_2_ but inhibit LH in the absence of sex steroids (10). It was therefore tentative to speculate that NKA could induce LH release through the stimulation of NKB given the absence of NK2R in Kiss1 and GnRH neurons. To test this hypothesis, first we co-administered NK2R and NK3R agonists to OVX + E_2_ WT mice and observed that the increase in LH was similar in groups injected with an individual dose of each agonist or the combination of both (**Figure 6 A**), discarding the possibility of an additive effect of NKA and NKB action on LH release and suggesting a possible common pathway. Next, to evaluate if NKA requires NKB signaling to induce LH release, the LH response to NK2R-Ag was tested in the presence of an NK3R antagonist (**Figure 6 B**) and in NKB deficient (*Tac2*KO) mice (**Figure 6 C**). In both cases, NK2R-Ag was able to significantly stimulate LH release indicating that NK2R activation induces LH release independently of the presence of NKB or its receptor NK3R. However, NK2R signaling requires the presence of kisspeptin as *Kiss1*KO mice replaced with E_2_ did not display any effect on LH release (WT Basal: 0.27 ± 0.06 ng/mL, WT NK2R-Ag: 0.57 ± 0.09 ng/mL; *p < 0.05; *Kiss1*KO Basal: 0.21 ± 0.02 ng/mL, *Kiss1*KO NK2R-Ag: 0.29 ± 0.04 ng/mL; not significant).

**6.**
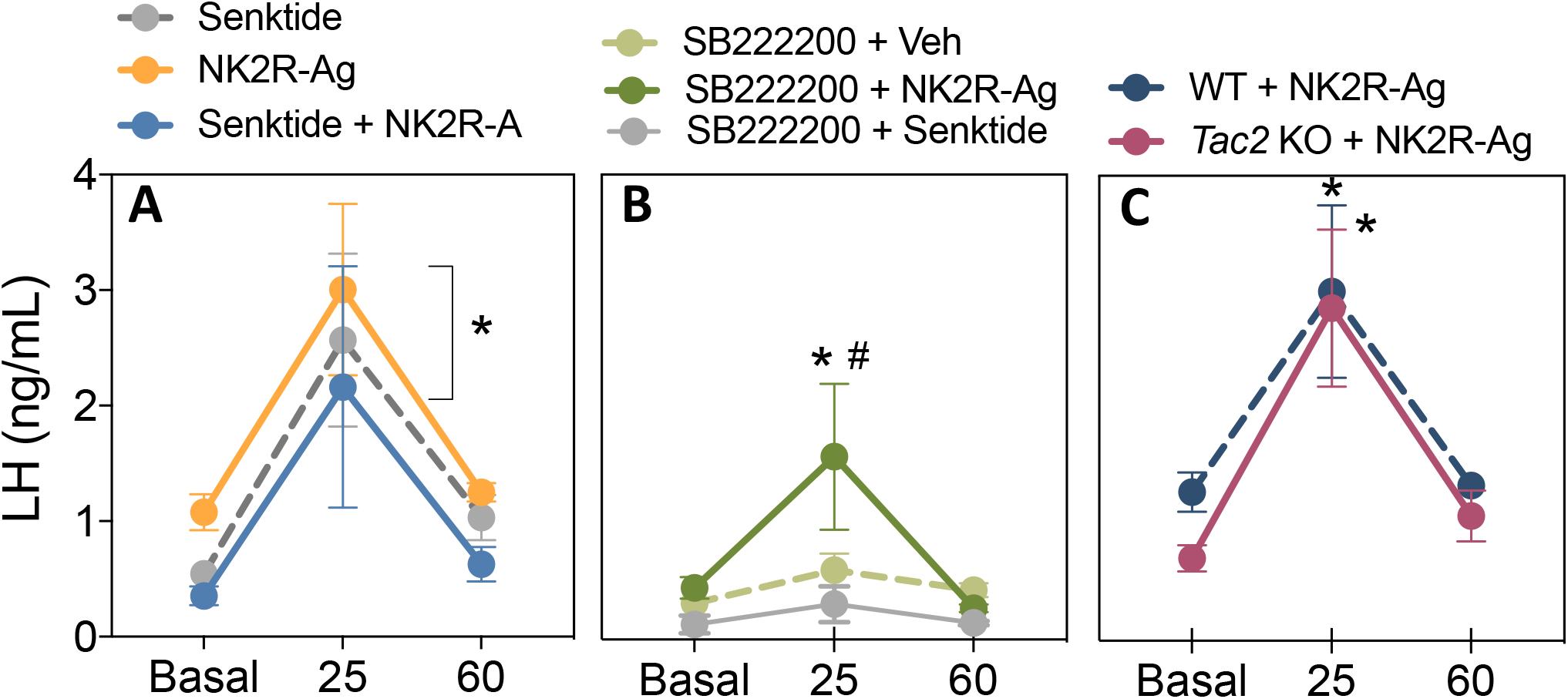
The stimulatory action of NKA is independent of NKB in the presence of physiological circulating levels of E2. **(A)** LH release before (basal), 25 and 60 min after the icv injection of NK2R-Ag, senktide or the co-administration of both (600 pmol/5ul/icv) in WT OVX+E_2_ females (n ≥ 5 per group). *p<0.05 vs. corresponding basal levels. (2 Way ANOVA followed by Tukey *post hoc* test). **(B)** LH levels before (basal) SB222200 (7 nmol5ul/icv) administration and at 25 and 60 minutes after injection of NK2R-Ag (600 pmol/5ul/icv), senktide (600 pmol/5ul/icv) or vehicle (0.9% NaCl/5ul/icv) in WT OVX+E_2_ females (n ≥ 4 per group). *p<0.05 vs. corresponding basal levels. # p<0.05 vs. corresponding control mice at the same time point (2 Way ANOVA followed by Tukey *post hoc* test). **(C)** LH before (Basal) and at 25 and 60 minutes after injection of NK2R-Ag (600 pmol/5ul/icv) in OVX+E_2_ WT and *Tac2*KO females (n≥5 per group). *p<0.05 vs. corresponding basal levels (2 Way ANOVA followed by Tukey *post hoc* test).

### 7. The inhibitory action of NKA is NKB and dynorphin dependent

In the next set of experiments, we sought to determine whether the inhibitory action of NKA/NK2R on LH release in the absence of sex steroids (i.e. OVX) is mediated by NKB and dynorphin. First, we showed that the inhibitory action of NK2R-Ag + senktide was similar to that of senktide alone, suggesting (as in the presence of E_2_) that there is no additive effect of both tachykinins in the inhibition of LH (**Figure 7 A**). The use of the specific NK3R antagonist already decreased LH in OVX animals, in line with recent literature showing that blockade of NK3R decreases LH pulsatility (22-25); however, co-administration of the NK3R antagonist and the NK2R-Ag failed to induce a further decrease in LH, suggesting that NK3R signaling is required for the inhibitory action of NKA in the absence of E_2_ (**Figure 7 B**). Moreover, as previously described in rats, NK2R signaling requires dynorphin to inhibit LH (26), which is abolished after the blockade of the kappa and mu receptor, KOR and MOR respectively, using naloxone (**Figure 7 C**). Of note, naloxone alone also inhibited LH release in OVX mice, in line with our previous reports in OVX *Pdyn*KO and *Oprk1*KO mice (dynorphin KO and KOR KO mice, respectively) (27), suggesting that the absence of the inhibitory signal of dynorphin leads to a significant decrease in the ability of the mouse to secrete LH, probably due to disruption of the LH pulse generator mechanism (28). Next, we assessed the action of NK2R-Ag in the absence of NKB (*Tac2*KO mice) to further confirm the data obtained after NK3R blockade. Unexpectedly, we observed that the absence of NKB leads to a robust induction of LH release, uncovering an action that is not present in WT OVX regardless of whether a functional NK3R is present or antagonized.

**7.**
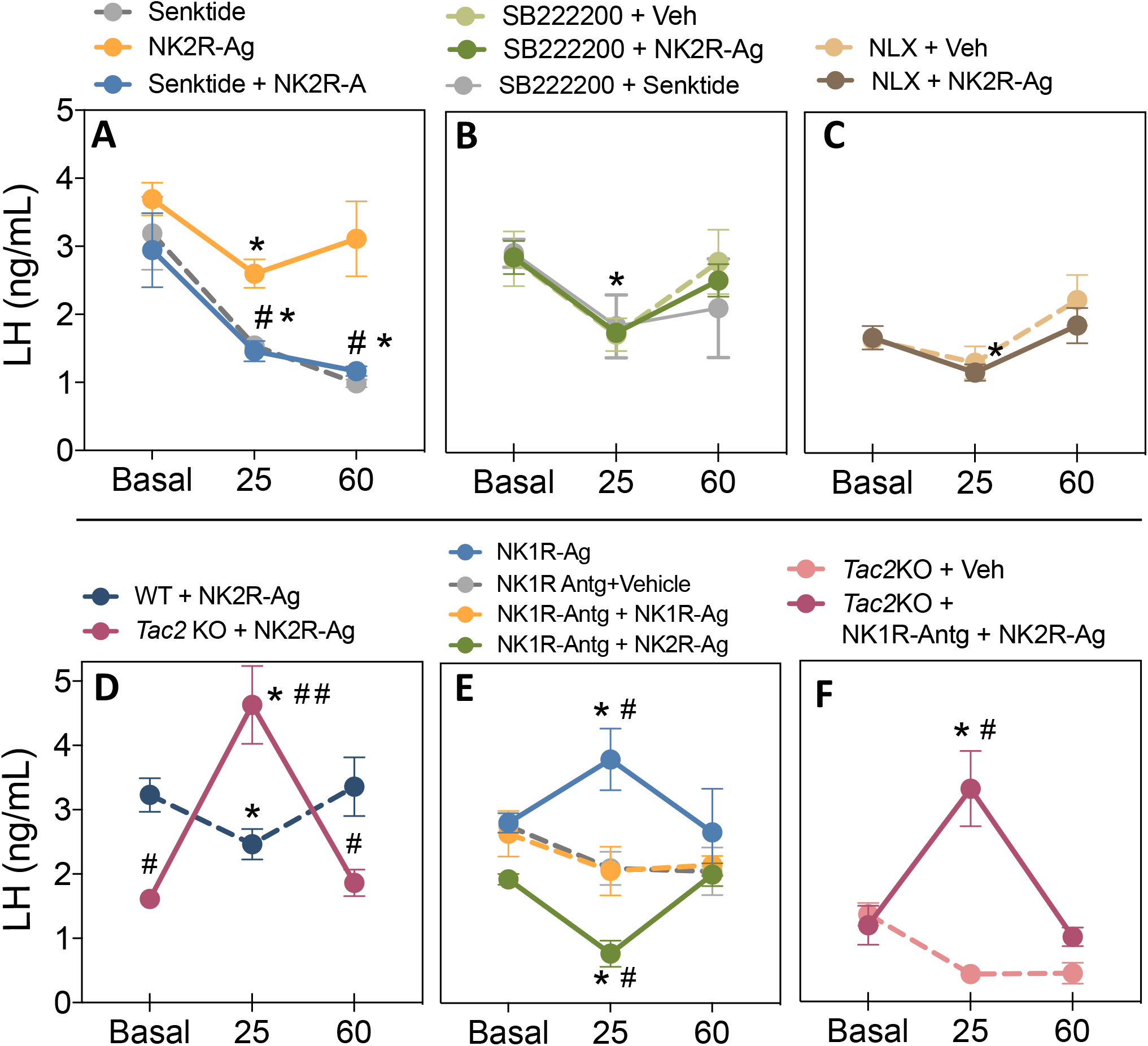
The inhibitory action of NKA is NKB and dynorphin dependent. **(A)** LH release before (basal), 25 and 60 min after icv injection of NK2R-Ag, senktide or the co-administration of both (600 pmol/5ul/icv) in WT OVX females (n≥5 per group). *p<0.05 vs. corresponding basal levels; # p<0.05 vs. NK2R-Ag at the same time point (2 Way ANOVA followed by Tukey post hoc test). **(B)** LH levels before (basal) SB222200 (7 nmol5ul/icv) injection and at 25 and 60 minutes after injection of NK2R-Ag (600 pmol/5ul/icv), senktide (600 pmol/5ul/icv) or vehicle (0.9% NaCl/5ul/icv) in WT OVX females (n≥5 per group). *p<0.05 vs. corresponding basal levels (2 Way ANOVA followed by Tukey *post hoc* test). **C)** LH levels before (basal), 25 and 60 minutes after injection of naloxone (5mg/kg/100ul/ip) or vehicle (0.9% NaCl/100ul/ip) in WT OVX females (n≥5 per group). *p<0.05 vs. corresponding basal levels (2 Way ANOVA followed by Tukey *post hoc* test). **(D)** LH levels before (basal) and at 25 and 60 minutes after injection of NK2R-Ag (600 pmol/5ul/icv) in OVX WT and OVX *Tac2*KO females (n≥5 per group). *p<0.05 vs. corresponding basal levels (2 Way ANOVA followed by Tukey *post hoc* test). # p<0.05; ## p<0.01 vs. corresponding WT levels (2 Way ANOVA followed by Tukey *post hoc* test). **E)** LH levels before (basal) and at 25 and 60 minutes after injection of NK1R-Ag (600 pmol/5ul/icv), NK2R-Ag (600 pmol/5ul/icv) or vehicle (0.9% NaCl/5ul/icv) in WT OVX females (n≥5 per group). Groups 2 – 4 were pre-injected 30 min before the basal sample with NK1R-Antg (2 nmol/5ul/icv). *p<0.05 vs. corresponding basal levels (2 Way ANOVA followed by Tukey post hoc test); # p<0.05 vs. NK1R-Antg + vehicle injected mice levels. **(F)** LH release before (basal) NK1R-Antg (2 nmol/5ul/icv) injection and at 25 and 60 minutes after injection of NK2R-Ag (600 pmol/5ul/icv) in *Tac2* KO OVX females (n≥5 per group). *p<0.05 vs. corresponding basal levels (2 Way ANOVA followed by Tukey post hoc test); # p<0.05 vs. vehicle injected mice.

Because we have observed that VMH Tac1 neurons co-express *Tacr2* (NK2R) (**Figure 4**), we hypothesized that NKA could also induce LH release through the stimulation of SP from Tac1 neurons that, in turn, would activate Kiss1 neurons (10). To test this hypothesis, we administered the NK2R-Ag in the presence of a NK1R antagonist (proven to efficiently block the action of a NK1R-agonist (**Figure 7, E**)) in WT and *Tac2*KO OVX mice. In both cases, NK2R-Ag was able to replicate the effect seen in WT (inhibition of LH, **Figure 7 E**) and *Tac2*KO mice (stimulation of LH, **Figure 7 F**). Lastly, we confirmed that this action is kisspeptin-dependent by showing complete absence of LH response to the administration of NK2R-Ag in *Kiss1* KO mice (WT: Basal 2,90 ± 0.42 ng/mL, NK1R-Ag 2.08 ± 0.32 ng/mL, *p < 0.05; *Kiss1*KO: Basal 0.32 ± 0.06 ng/mL, NK1R-Ag 0.30 ± 0.02 ng/mL, not significant).

## Discussion

In the present study, we show that all tachykinins are essential for the normal timing of puberty onset in both sexes and critical for the maintenance of fertility in females. Humans and mice with deficient NKB/NK3R signaling present delayed puberty onset (4, 5, 7, 8). We have also described previously that SP is critical for the proper timing of puberty onset in the mouse (16, 17) and here we extend that finding to include NKA as well.

The reproductive phenotype of *TAC3/TACR3* deficient patients (without NKB/NK3R signaling) has spurred controversy as their infertile phenotype has been shown to reverse spontaneously in some patients leading to successful pregnancies (6). This phenotype resembled the fertile phenotype of both *Tac2* and *Tacr3* KO mice (7, 8). The current model of NKB’s action proposes that NKB plays a role before puberty by stimulating the release of kisspeptin (29) and that after puberty it participates in the shaping of kisspeptin-GnRH-LH pulses as part of the GnRH pulse generator within the arcuate/infundibular nucleus (30). Therefore, the explanation for the reversal of the HH phenotype in patients and mice without this seemingly critical factor for GnRH pulsatility remained unknown and demanded further investigation. Because tachykinins have been described to cross-activate all three NK receptors (31, 32), we hypothesized that in the absence of NKB signaling, SP and/or NKA could compensate for the role of NKB and eventually be able to activate the gonadotropic axis. Our present results support this hypothesis, as the complete absence of all tachykinins in double *Tac1/Tac2* KO mice *1)* recapitulates the delay in puberty onset observed in NKA/SP and NKB null models (8, 16), and *2)* leads to severe reproductive defects (80% of infertile mice) in females in striking difference with the reproductive phenotype of each studied tachykinin KO model separately (i.e. *Tac1*KO, *Tac2*KO and *Tacr3*KO), which displayed 100% of successful pregnancies in all cases (7, 16). However, although *Tac1/Tac2* KO males presented delayed puberty onset, as documented previously for *Tac1*KO mice (17), all of them were able to father litters. The sex difference in the fertility phenotype is remarkable and points to the fact that tachykinins may be important for the timing of puberty onset in both sexes and proper pulsatile release of LH in adult females, while additional (yet unknown) mechanisms are able to maintain proper activation of the gonadotropic axis in adult males. It is also possible that in the female, tachykinins act at the level of Kiss1 neurons in the anteroventral periventricular nucleus (AVPV/PeN), exclusive to the female brain and involved in the positive feedback of sex steroids that leads to ovulation (33). In that case, the absence of tachykinin action in these neurons could prevent them from mounting a proper ovulatory response, as suggested by their lack of corpora lutea and almost complete infertility of *Tac1/Tac2* KO females in this study, leading to the sexual differentiation in the infertile vs. fertile phenotype of females and males, respectively. Supporting this contention, we have previously described that a fraction of Kiss1 neurons in the ARC and AVPV/PeN express NK1R and NK3R mRNA, but not NK2R (10). In this study, we show that the NK2R transcript (*Tacr2*) is present in additional (unknown) neurons of the ARC and in Tac1 neurons of the VMH, suggesting the existence of autosynaptic loops in these neurons similar to NKB’s action in ARC Kiss1 neurons, which could serve as an indirect mechanism to modulate Kiss1 neurons through the regulation of SP release.

Our present findings also demonstrate that TACs are involved in the rapid response of Kiss1 neurons to the absence of negative feedback, i.e. increase in LH release after gonadectomy. Tachykinin deficient mice (i.e. *Tac1/Tac2* KO) showed a significant delay in the post-gonadectomy rise of LH levels 48h after surgery. However, LH levels are restored to normal values by 7d post-surgery, indicating that tachykinins are not necessary for the activation of Kiss1 neurons in the mid-long term but contribute to the rapid response of the animal to changes in the sex steroid milieu. Interestingly, in both sexes, the acute administration of kisspeptin or GnRH led to a diminished response of LH in *Tac1*KO mice (as we described previously in male mice (17)) and in double *Tac1/Tac2*KO mice. While this may reflect a defect at the level of the pituitary, both genotypes were able to reach normal compensatory rises of LH after gonadectomy, as indicated above, supporting the ability to mount a proper kisspeptin-GnRH-LH response and the presence of normal gonadotrope function. This is in line with the delay in the response to GDX described above, indicating the difficulty of these animals to mount a rapid response to GnRH and suggesting that substance P may facilitate the rapid response of gonadotropes to GnRH at the pituitary level, in agreement with previous *in vitro* studies (34-37).

In this study we also addressed the question of whether the analogous action of NKA and NKB in the regulation of LH release (i.e. stimulation in the presence of E_2_ and inhibition in its absence (10)) is due to a converging mechanism of action of NKA onto NKB signaling. Despite these similarities, our data using NK3R antagonists and *Tac2*KO mice after OVX and E_2_ replacement, clearly demonstrate that the stimulatory action of NKA on LH is NKB *independent* but kisspeptin-dependent, suggesting the existence of a yet unknown population of NKA-responsive neurons that in turn activate Kiss1 neurons to induce kisspeptin/GnRH release. On the contrary, NKA had been shown to inhibit LH via dynorphin in the rat (19, 26), in a similar mechanism to that described for NKB in the absence of E_2_ (38). Here, we further demonstrate that this inhibitory action of NKA is mediated by the activation of the NKB–dynorphin pathway because the blockade of both NK3R and KOR receptors ablates the inhibitory action of NKA. However, we unexpectedly observed that in the absence of NKB, NKA no longer inhibits LH in the absence of E_2_ but rather significantly stimulates it (our present data in *Tac2*KO mice) in a process that is also SP-independent. This suggests that when NKB is present, its inhibitory action (after being induced by NKA) is downstream of any stimulation induced by NKA which, as documented by our ISH data, would act upstream of Kiss1 neurons. However, in the absence of NKB (and in the presence of a functional NK3R, i.e. in *Tac2*KO mice), NKA is able to further stimulate LH release through a mechanism that requires the activation of NK3R (because it is absent when this receptor is blocked). This demonstrates that NKA’s action on LH release is inherently stimulatory and requires a functional NK3R, but not NKB.

Overall, in this study we have demonstrated that tachykinins are essential for the proper timing of puberty onset in males and females and fertility maintenance in females. The absence of a single tachykinin system may induce reproductive defects that can range from subtle to severe but can eventually be reversed, and lead to successful reproduction, as observed in humans and mice lacking NKB signaling. However, under the total absence of tachykinins, reproductive function is severely compromised in the female supporting a compensatory role of the other tachykinin systems, i.e. SP/NKA, in maintaining fertility. Despite this significant compensatory effect in the tachykinin systems, it is remarkable that 20% of female *Tac1/Tac2*KO mice were able to deliver pups. This suggests that higher hierarchical regulatory systems of Kiss1 neurons, e.g. tachykinins, are not indispensable and additional mechanisms may develop to achieve reproductive success although at very low levels. Male mice however, retain reproductive capabilities after complete tachykinin removal, suggesting the existence of additional mechanisms that ensure sufficient LH pulsatile release and/or the need of proper tachykinin signaling in females to induce ovulation. This study offers new insights into the interaction and mechanism of action of tachykinins in the control of LH release, especially related to NKA-NKB interaction, which remained largely unexplored, and challenges the existing notion of NKB (and tachykinins in general) as critical regulators of the GnRH pulse generator, at least in males. This redundant tachykinin systems and the ability of NKA to compensate for NKB’s action uncovers a novel mechanism with potential clinical application to activate the gonadotropic axis in *TAC3-*deficient (but not *TAC3R*-deficient) patients, through the exogenous administration of NKA or NK2R agonists to achieve reproductive success as occurs in sporadic HH reversal patients.

## Materials and Methods

### Mice

Wild-type (WT) female C57Bl/6 mice were purchased from Charles River Laboratories International, Inc. *Tac2* KO (knockout, KO) mice were obtained from Dr. Seminara (MGH) (8). *Tac1/Tac2* KO were generated by crossing *Tac1*KO (The Jackson Laboratories, stock No. 004103) and *Tac2*KO mice. All animal studies were approved by the Harvard Medical Area Standing Committee on the Use of Animals in Research and Teaching in the Harvard Medical School Center for Animal Resources and Comparative Medicine. Mice were maintained in a 12:12 h light/dark cycle and were fed standard rodent chow diet and water ad libitum. Genotyping was conducted by PCR analyses on isolated genomic DNA from tail biopsies.

### Reagents

The agonists of NK1R (GR73632), NK2R (GR64349) and NK3R (senktide), and the antagonists of NK3R (SB 222200) and NK1R (RP67580) were purchased from Tocris. Naloxone Hydrochloride (opioid receptor antagonist) and GnRH, were purchased from Sigma Aldrich. Mouse kisspeptin-10 (Kp-10) was purchased from Phoenix pharmaceutical. All drugs were dissolved in saline (0.9% NaCl), except for SB 222200 and RP67580, which were dissolved in 5% DMSO. Saline (0.9% NaCl) was used as vehicle in all of our experiments. Doses and timings for hormonal analyses were selected on the basis of previous studies (10, 21, 39).

### Experimental design

#### General procedures

For intracerebroventricular (icv) injection, 2-3 days before the experiment, the mice were briefly anesthetized with isoflurance and a small hole was bored in the skull 1 mm lateral and 0.5 mm posterior to bregma with a Hamilton syringe attached to a 27-gauge needle fitted with polyethylene tubing, leaving 3.5 mm of the needle tip exposed. Once the initial hole was made, all subsequent injections were made at the same site. For icv injections, mice were anesthetized with isoflurane for a total of 2–3 min, during which time 5 μl of solution were slowly and continuously injected into the lateral ventricle. The needle remained inserted for approximately 60 sec after the injection to minimize backflow up the needle track. Mice typically recovered from the anesthesia within 3 min after the injection. For hormonal analyses, blood samples (4 μl) were obtained from the tail and stored at −80°C until hormonal determination. The dose and time of collection were selected based on our previous studies (10).

#### Study 1: Reproductive maturation and function of Tac1/Tac2 KO female mice

Reproductive phenotyping (presence of vaginal opening (VO), timing of first estrus, ovarian weights) and fertility parameters (percentage of pregnancies, litter size and time to the first litter) were measured in female mice.

Littermate WT (n=12), *Tac1* KO (n=12), *Tac2* KO (n=5) and *Tac1/Tac2* KO (n=8) females were monitored daily from p25 (postnatal day) for: body weight (BW) and progression to VO (as indicated by complete canalization of the vagina). In addition, WT and *Tac1/Tac2* KO females were monitored for the timing of first estrus during 90 days after the day of VO (first day with cornified cells determined by daily [in the morning] vaginal cytology). In addition, estrous cyclicity was monitored by daily vaginal cytology, for a period of 30 in WT and *Tac1/Tac2* KO (n≥8). Cytology samples were obtained every morning (10 am) and placed on a glass slide for determination of the estrous cycle under the microscope as previously described (40).

The fertility assessment was performed by breeding adult WT (n=3) or *Tac1/Tac2* KO (n=10) females with WT male previously proven to father litters. The time to the first litter and number of pups per litter were monitored.

The ovarian ultra-structure was analyzed in adult (3–4 month old) mice of the two genotypes: WT and *Tac1/Tac2* KO (n=4). Ovaries were collected, weighed and fixed in Bouin’s solution. The tissues were embedded in paraffin and sectioned (10 μm) for hematoxylin and eosin staining (Harvard Medical School Rodent Pathology Core) and images acquired under ×4 magnification. The ovaries were analyzed for presence of corpora lutea (CLs) per section. Each value represents the number of CLs of 1 representative section from the middle line of one ovary per animal.

#### Study 2: Reproductive maturation and function of the Tac1/Tac2 KO male mouse

Prepubertal littermate WT (n=11), *Tac1* KO (n=6), *Tac2* KO (n=6) and *Tac1/Tac2* KO (n=14) males were monitored daily from p25 for preputial separation as an indirect marker of puberty onset, and body weight was measured at the average age of puberty onset (PND28).

Adult WT (n=3) or *Tac1/Tac2* KO (n=7) littermate mice (>PND75) were placed with proven fertile WT females. Time to delivery and numbers of pups were monitored.

The testes’ ultra-structure was analyzed in adult (3–4 month old) mice of the two genotypes: WT and *Tac1/Tac2* KO (n=4). Testes were collected and processed as described above for ovaries.

#### Study 3: Characterization of the postgonadectomy response of LH in females

LH levels were measured in intact (diestrus in the morning) WT (n=9), *Tac2* KO (n=4) and *Tac1/Tac2* KO (n=6) adult (3-4 month) females and compared with 2d and 7d (days) after bilateral ovariectomy (OVX). In addition, we assessed the pulsatile secretion of LH in adult OVX females (4-week after OVX) *Tac1/Tac2* KO mice and WT littermates (n=6-7 per group). Mice were handled daily to allow acclimation to sampling conditions for three weeks prior to the experiment. Pulsatile measurements of LH secretion were assessed by repeated blood collection through a single excision at the tip of the tail, as described previously (41). The tail was cleaned with saline and then massaged to take a 4 ul blood sample with a pipette. Whole blood was immediately diluted in 116 ul of 0.05% PBST, vortexed, and frozen on dry ice. Samples were stored at −80°C for a subsequent LH ELISA. We collected 16 sequential blood samples over a 90 minutes sampling period.

#### Study 4: Effect of Kp-10 and GnRH administration on LH release in adult female mice

Hormonal (LH) responses to known stimulators of GnRH and/or gonadotropin secretion were studied in WT (n=7), *Tac1* KO (n=8), *Tac2* KO (n=5) and *Tac1/Tac2* KO (n=7) mice. The mice were injected with GnRH (0.25 ug/100μl/ip) and kisspeptin-10 (Kp-10) (50 pmol/mouse in 5 μl/icv), and blood samples were obtained at 15-min after icv administration of Kp-10 and 30 min after ip administration of GnRH. Doses and routes of administration were selected in the basis of previous references (42, 43).

#### Study 5: Expression of Tacr1,Tacr2, Tacr3, Kiss1 and Pdyn in the mediobasal hypothalamus (MBH) of female mice

We aimed to determine if there are changes in the expression of *Tacr1, Tacr2, Tacr3, Kiss1*, and *Pdyn* in the mediobasal hypothalamus (MBH), the site that includes the arcuate nucleus (ARC) between WT and *Tac1/Tac2* KO intact females.

Total RNA from the MBH was isolated using TRIzol reagent (Invitrogen) followed by chloroform/isopropanol extraction. RNA was quantified using NanoDrop 2000 spectrophotometer (Thermo Scientific), and 1 μm of RNA was reverse transcribed using iScript cDNA synthesis kit (Bio-Rad). Quantitative real-time PCR assays were performed on an ABI Prism 7000 sequence detection system, and analyzed using ABI Prism 7000 SDS software (Applied Biosystems). The cycling conditions were the following: 2 min incubation at 95°C (hot start), 45 amplification cycles (95°C for 30 s, 60°C for 30 s, and 45 s at 75°C, with fluorescence detection at the end of each cycle), followed by melting curve of the amplified products obtained by ramped increase of the temperature from 55 to 95°C to confirm the presence of single amplification product per reaction. For data analysis, relative standard curves were constructed from serial dilutions of one reference sample cDNA and the input value of the target gene was standardized to *Hprt* levels in each sample. The primers used are listed in **Table 1**.

**Table:**
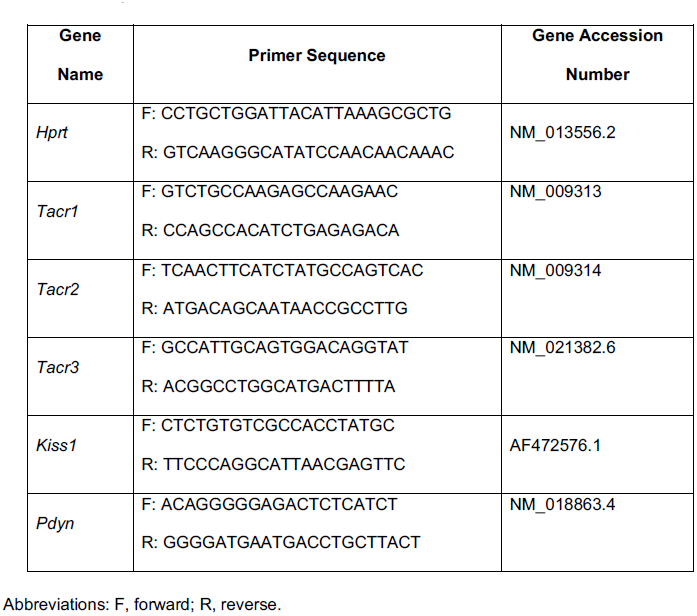
List of primers used for PCR determination.

#### Study 6: Characterization of the postgonadectomy response of LH in males

Bilateral removal of testes from 3-4-month-old males was performed with light isoflurane anesthesia. Briefly, the ventral skin was shaved and cleaned to perform one small incision in the skin and abdominal musculature of the abdomen. Once the gonads were identified and excised, the muscle incision was sutured and the skin was closed with surgical clips. LH levels were measured in WT (n=13), *Tac2* KO (n=4) and *Tac1/Tac2* KO (n=10) mice. Blood samples were collected before and 2d and 7d after bilateral gonadectomy (GNX).

#### Study 7: Effect of Kp-10 and GnRH administration on LH release in adult male mice

Hormonal (LH) responses to known stimulators of GnRH and/or gonadotropin secretion were studied in WT (n=7-20), *Tac1* KO (n=4-7), *Tac2* KO (n=8-12) and *Tac1/Tac2* KO (n=8-18) mice. The mice were injected with GnRH (0.25 ug/100μl/ip) and kisspeptin-10 (Kp-10) (50 pmol/mouse in 5 μl/icv), and blood samples were obtained at 15-min after icv administration of Kp-10 and 30 min after ip administration of GnRH. Doses and routes of administration were selected in the basis of previous references.

#### Study 8: Effect of chronic administration of NK2R-Ag in pre-pubertal mice

To address if NKA/NK2R signaling plays a role in puberty onset, in the next experiment, we performed a systemic chronic (p23 – 32) administration of NK2R-Ag (3 nmol/100μl/ ip) or vehicle (0.9% NaCl) to female mice (n≥6 per group). Reproductive maturation (progression of VO) was monitored daily. BW was registered at day 30 when 50% of control females showed VO. Also, uterus and ovarian weight, as well as serum LH were determined at day 32, last day of NK2R-Ag administration.

#### Study 9: Interaction between NKA, NKB and SP in the stimulation of LH release in the presence of estradiol

In this study, adult female mice were subjected to bilateral OVX via abdominal incision under light isofluorane anesthesia 1 week before hormonal tests. Immediately after OVX, capsules filled with E_2_ or vehicle (sesame seed oil) were implanted subcutaneously (sc) via a small midscapular incision at the base of the neck; wound clips were used to close the incision. Silastic tubing (15 mm long, 0.078 in inner diameter, 0.125 in outer diameter; Dow Corning) was used for capsule preparation. Dilutions of crystalline E_2_ at a low dose (50 μg/mL, in sesame oil) were used to fill capsules. After capsules were filled, the end of the capsule was sealed with silicone cement and allowed to cure overnight. On the day before surgery, implants were washed twice for 10 min in changes of 100% ethanol and then placed in sterile physiological saline overnight.

First, we analyzed the possible additive effect of NK2R-Ag and senktide in LH secretion in the presence of sex steroids. The LH level was measured at 25 and 60 min after icv injection of NK2R-Ag, senktide or the co-administration of both drugs. Then, OVX+E_2_ females (n≥5 per group) were pretreated with SB222200 (7 nmol), 60 minutes prior to the icv injection of NK2R-Ag (600 pmol, Senktinde or vehicle (0.9% NaCl). Blood samples were collected before SB222200 injection (Basal) and at 25 and 60 minutes after injection. Additionally, we further analyzed the action of NK2R-Ag in the absence of NKB signaling using *Tac2* KO OVX+E_2_ females (n≥5 per group) that were injected with NK2R-Ag (600 pmol) and blood samples were collected before and 25 and 60 min after injection. Finally, in order to evaluate the whether the action of NKA requires kisspeptin to stimulate LH release, we used *Kiss1* KO OVX+E_2_ females (n≥5 per group) and LH levels were measured 25 min after icv injection of NK2R-Ag (600 pmol).

#### Study 10: Interaction between NKA, NKB and SP in the inhibition of LH release in ovariectomized (OVX) female mice

Adult females were subjected to bilateral OVX via abdominal incision under light isofluorane anesthesia 1 week before hormonal tests. First, we analyzed the possible additive effect of NK2R-Ag and senktide in the inhibition of LH secretion in the absence of sex steroids. LH level was measured at 25 and 60 min after icv injection of NK2R-Ag, senktide or the co-administration of both drugs. In the next experiment we intended to assess the role of NKB in the inhibition of LH secretion achieved by NKA in OVX females. To this end, LH responses to NK2R-Ag were evaluated after blockade of the effects of NKB using SB222200 (7 nmol) as a selective antagonist of NK3R. For this purpose, adult OVX female mice were pretreated with SB222200, 60 minutes prior to the icv injection of NK2R-Ag (600 pmol). Blood samples were collected before SB222200 injection (Basal) and at 25 and 60 minutes after vehicle or NK2R-Ag injection. In addition, we analyzed the role of endogenous opioids in the control of adult OVX females LH secretion and in the modulation of responses to NK2R-Ag. To this end, LH responses to NK2R-Ag were measured after the blockade of the opioid receptors κ and μ (KOR and MOR) using naloxone (5mg/kg/100ul/ip). The animals were injected with naloxone, 12 hours and 60 minutes prior to the icv injection of NK2R-Ag (600 pmol). Blood samples were collected before naloxone injection and at 25 and 60 minutes after NK2R-Ag injection. We further analyzed the action of NK2R-Ag in the absence of NKB signaling and sex steroids using *Tac2* KO OVX females (n≥5 per group), which were injected with NK2R-Ag (600 pmol) and blood samples were collected before and 25 and 60 min after injection. In addition, we evaluated the role of SP in the action of NK2R-Ag action in the inhibition of LH in WT OVX and *Tac2*KO OVX mice. WT OVX and *Tac2*KO OVX females (n≥5 per group) were injected with NK1R-Antg (2 nmol), 30 min previously to the NK2R-Ag (600 pmol), NK1R-Ag (600 pmol) or vehicle administration. Blood samples were collected before and 25 and 60 min after injection. Finally, we assess the ability of NK2R signaling to modulate LH release in the absence of sex steroids and kisspeptin using Kiss1 KO OVX females (n≥5 per group) were injected with NK2R-Ag (600 pmol) and LH levels were measured at 25 and 60 min after icv injection.

#### In situ hybridization (ISH)

To determine the presence of co-expression between *Tac2r* and *Tac1* mRNA in key areas (ventromedial nucleus, VMN; and ARC) dual fluorescence ISH was performed in additional tissue samples from OVX+sham and OVX+E_2_ animals. We used probes for *Tac2r*-C1 and *Tac1*-C2 obtained from ACDBio and used the RNAscope method per their protocol (ACDBio). The brains were removed for ISH, fresh frozen on dry ice, and then stored at −80°C until sectioned. Five sets of 20-μm sections in the coronal plane were cut on a cryostat, from the diagonal band of Broca to the mammillary bodies, thaw mounted onto SuperFrost Plus slides (VWR Scientific), and stored at −80°C. A single set was used for ISH experiment (adjacent sections 100 μm apart).

#### Hormone measurements

LH was measured by a sensitive sandwich ELISA for the assessment of whole blood LH concentrations. A 96-wellhigh-affinity binding microplate (9018; Corning) was coated with 50uL of capture antibody (monoclonal antibody, anti-bovine LH beta subunit, 518B7; University of California) at a final dilution of 1:1000 (in 1XPBS, 1.09 g of Na2HPO4 [an-hydrous], 0.32 g of NaH2PO4 [anhydrous], and 9g of NaCl in1000 mL of distilled water) and incubated overnight at 4°C. To minimize unspecific binding of the capture antibody, wells were incubated with 200uL of blocking buffer (5% [w/v] skim milk powder in 1XPBS-T (1XPBS with 0.05% Tween20) for 2hours at room temperature (RT).A standard curve was generated using a 2-fold serial dilution of LH (reference preparation, AFP-5306A; National Institute of Diabetes and Digestive and Kidney Diseases National Hormone and Pituitary Program [NIDDK-NHPP]) in 0.2% (w/v) BSA-1XPBS-T. The LH standards and blood samples were incubated with 50 uL of detection antibody (polyclonal antibody, rabbit LH antiserum, AFP240580Rb; NIDDK-NHPP) at a final dilution of 1:10000 for 1.5 hours (at RT). Each well containing bound substrate was incubated with 50 ul of horseradish peroxidase conjugated antibody (poly-clonalgoatanti-rabbit, D048701– 2; DakoCytomation) at a final dilution of 1:2000. After a 1.5-hour incubation, 100Ul of o-phenylenediamine (002003;Invitrogen), substrate containing 0.1% H2O2 was added to each well and left at RT for 30minutes. The reaction was stopped by addition of 50 uL of 3M HCl to each well, and absorbance of each well was read at a wave length of 490 nm (Sunrise; Tecan Group). The concentration of LH in whole blood samples was determined by interpolating the OD values of unknowns against a nonlinear regression of the LH standard curve (41).

#### Statistical Analysis

All data are expressed as the mean ± SEM for each group. A two tailed unpaired t-Student test or a one- or two-way ANOVA test followed by Tukey or Newman Kleus *post-hoc* test was used to assess variation among experimental groups. Significance level was set at P < 0.05. All analyses were performed with GraphPad Prism Software, Inc (San Diego, CA).

#### Statistical Analysis of LH pulses

Mice LH concentration time series were analyzed using a MATLAB-bases algorithm. It is a for loop written in the code to determine which LH peaks are considered pulses. This for loop states that any peak whose height is 20% greater than the heights of the 2 previous peaks as well as 10% greater than the height of the following peak is considered a pulse. There is also a condition written into the code that is specific for the second time interval (i=2) that states that the peak at the second-time interval only needs to be 20% greater than the single peak that comes before it to be considered a pulse.

#### Study Approval

All animal care and experimental procedures were approved by the National Institute of Health, and Brigham and Women’s Hospital Institutional Animal Care and Use Committee, protocol #05165. The Brigham and Women’s Hospital is a registered research facility with the U.S. Department of Agriculture (#14-19), is accredited by the American Association for the Accreditation of Laboratory Animal Care, and meets the National Institutes of Health standards as set forth in the Guide for the Care and Use of Laboratory Animals (DHHS Publication No. (NIH) 85-23 Revised 1985).

## Author contributions

SL and VMN conceived and designed the research. SL, CF, RT, SS, CAM and AG conducted experiments. SBS provided critical mouse lines. SL and VMN contributed to data analysis. SL and VMN wrote the manuscript, and all authors contributed to manuscript editing.

## Acknowledgments

This work was supported by R01 HD090151 and Women’s Brain Initiative of the Brigham and Women’s Hospital to V.M.N. and The International Brain Research Organization (IRBO). Research Fellowship to RT. The authors declare no competing financial interests

## References

1. Lasaga M, Debeljuk L. Tachykinins and the hypothalamo-pituitary-gonadal axis: An update. Peptides. 2011;32(9):1972–8.

2. Cao YQ, Mantyh PW, Carlson EJ, Gillespie AM, Epstein CJ, Basbaum AI. Primary afferent tachykinins are required to experience moderate to intense pain. Nature. 1998;392(6674):390–4.

3. Fergani C, Navarro VM. Expanding the Role of Tachykinins in the Neuroendocrine Control of Reproduction. Reproduction (Cambridge, England). 2016;153(1):R1–R14.

4. Topaloglu AK, Reimann F, Guclu M, Yalin AS, Kotan LD, Porter KM, et al. TAC3 and TACR3 mutations in familial hypogonadotropic hypogonadism reveal a key role for Neurokinin B in the central control of reproduction. Nat Genet. 2009;41(3):354–8.

5. Young J, Bouligand J, Francou B, Raffin-Sanson ML, Gaillez S, Jeanpierre M, et al. TAC3 and TACR3 defects cause hypothalamic congenital hypogonadotropic hypogonadism in humans. J Clin Endocrinol Metab. 2010;95(5):2287–95.

6. Gianetti E, Tusset C, Noel SD, Au MG, Dwyer AA, Hughes VA, et al. TAC3/TACR3 mutations reveal preferential activation of gonadotropin-releasing hormone release by neurokinin B in neonatal life followed by reversal in adulthood. J Clin Endocrinol Metab. 2010;95(6):2857–67.

7. Yang JJ, Caligioni CS, Chan YM, Seminara SB. Uncovering novel reproductive defects in neurokinin B receptor null mice: closing the gap between mice and men. Endocrinology. 2012;153(3):1498–508.

8. True C, Nasrin Alam S, Cox K, Chan YM, Seminara S. Neurokinin B is critical for normal timing of sexual maturation but dispensable for adult reproductive function in female mice. Endocrinology. 2015:en20141862.

9. de Croft S, Boehm U, Herbison AE. Neurokinin B activates arcuate kisspeptin neurons through multiple tachykinin receptors in the male mouse. Endocrinology. 2013;154(8):2750–60.

10. Navarro VM, Bosch MA, Leon S, Simavli S, True C, Pinilla L, et al. The integrated hypothalamic tachykinin-kisspeptin system as a central coordinator for reproduction. Endocrinology. 2015;156(2):627–37.

11. Arisawa M, De Palatis L, Ho R, Snyder GD, Yu WH, Pan G, et al. Stimulatory role of substance P on gonadotropin release in ovariectomized rats. Neuroendocrinology. 1990;51(5):523–9.

12. Coiro V, Volpi R, Capretti L, Caiazza A, Marcato A, Bocchi R, et al. Luteinizing hormone response to an intravenous infusion of substance P in normal men. Metabolism: clinical and experimental. 1992;41(7):689–91.

13. Traczyk WZ, Pau KY, Kaynard AH, Spies HG. Modulatory role of substance P on gonadotropin and prolactin secretion in the rabbit. J Physiol Pharmacol. 1992;43(3):279–97.

14. Rance NE, Bruce TR. Neurokinin B gene expression is increased in the arcuate nucleus of ovariectomized rats. Neuroendocrinology. 1994;60(4):337–45.

15. Hrabovszky E, Borsay BA, Racz K, Herczeg L, Ciofi P, Bloom SR, et al. Substance p immunoreactivity exhibits frequent colocalization with kisspeptin and neurokinin B in the human infundibular region. PloS one. 2013;8(8):e72369.

16. Simavli S, Thompson IR, Maguire CA, Gill JC, Carroll RS, Wolfe A, et al. Substance p regulates puberty onset and fertility in the female mouse. Endocrinology. 2015;156(6):2313–22.

17. Maguire CA, Song YB, Wu M, Leon S, Carroll RS, Alreja M, et al. Tac1 Signaling is Required for Sexual Maturation and Responsiveness of GnRH Neurons to Kisspeptin in the Male Mouse. Endocrinology. 2017.

18. Ruiz-Pino F, Garcia-Galiano D, Manfredi-Lozano M, Leon S, Sanchez-Garrido MA, Roa J, et al. Effects and interactions of tachykinins and dynorphin on FSH and LH secretion in developing and adult rats. Endocrinology. 2015;156(2):576–88.

19. Sahu A, Kalra SP. Effects of tachykinins on luteinizing hormone release in female rats: potent inhibitory action of neuropeptide K. Endocrinology. 1992;130(3):1571–7.

20. Steinhoff MS, von Mentzer B, Geppetti P, Pothoulakis C, Bunnett NW. Tachykinins and their receptors: contributions to physiological control and the mechanisms of disease. Physiological reviews. 2014;94(1):265–301.

21. Navarro VM, Ruiz-Pino F, Sanchez-Garrido MA, Garcia-Galiano D, Hobbs SJ, Manfredi-Lozano M, et al. Role of neurokinin B in the control of female puberty and its modulation by metabolic status. The Journal of neuroscience: the official journal of the Society for Neuroscience. 2012;32(7):2388–97.

22. George JT, Kakkar R, Marshall J, Scott ML, Finkelman RD, Ho TW, et al. Neurokinin B Receptor Antagonism in Women With Polycystic Ovary Syndrome: A Randomized, Placebo-Controlled Trial. J Clin Endocrinol Metab. 2016;101(11):4313–21.

23. Li SY, Li XF, Hu MH, Shao B, Poston L, Lightman SL, et al. Neurokinin B receptor antagonism decreases luteinising hormone pulse frequency and amplitude and delays puberty onset in the female rat. Journal of neuroendocrinology. 2014;26(8):521–7.

24. Nakamura S, Wakabayashi Y, Yamamura T, Ohkura S, Matsuyama S. A neurokinin 3 receptor-selective agonist accelerates pulsatile luteinizing hormone secretion in lactating cattle. Biology of reproduction. 2017;97(1):81–90.

25. Noritake K, Matsuoka T, Ohsawa T, Shimomura K, Sanbuissho A, Uenoyama Y, et al. Involvement of neurokinin receptors in the control of pulsatile luteinizing hormone secretion in rats. The Journal of reproduction and development. 2011;57(3):409–15.

26. Kalra PS, Sahu A, Bonavera JJ, Kalra SP. Diverse effects of tachykinins on luteinizing hormone release in male rats: mechanism of action. Endocrinology. 1992;131(3):1195–201.

27. Navarro VM, Gottsch ML, Chavkin C, Okamura H, Clifton DK, Steiner RA. Regulation of gonadotropin-releasing hormone secretion by kisspeptin/dynorphin/neurokinin B neurons in the arcuate nucleus of the mouse. The Journal of neuroscience: the official journal of the Society for Neuroscience. 2009;29(38):11859–66.

28. Navarro VM. New insights into the control of pulsatile GnRH release: the role of Kiss1/neurokinin B neurons. Frontiers in endocrinology. 2012;3:48.

29. Gill JC, Navarro VM, Kwong C, Noel SD, Martin C, Xu S, et al. Increased neurokinin B (Tac2) expression in the mouse arcuate nucleus is an early marker of pubertal onset with differential sensitivity to sex steroid-negative feedback than Kiss1. Endocrinology. 2012;153(10):4883–93.

30. Clarkson J, Han SY, Piet R, McLennan T, Kane GM, Ng J, et al. Definition of the hypothalamic GnRH pulse generator in mice. Proceedings of the National Academy of Sciences of the United States of America. 2017;114(47):E10216–E23.

31. Deal MJ, Hagan RM, Ireland SJ, Jordan CC, McElroy AB, Porter B, et al. Conformationally constrained tachykinin analogues: potent and highly selective neurokinin NK-2 receptor agonists. Journal of medicinal chemistry. 1992;35(22):4195–204.

32. Seabrook GR, Bowery BJ, Hill RG. Pharmacology of tachykinin receptors on neurones in the ventral tegmental area of rat brain slices. Eur J Pharmacol. 1995;273(1-2):113–9.

33. Smith JT, Cunningham MJ, Rissman EF, Clifton DK, Steiner RA. Regulation of Kiss1 gene expression in the brain of the female mouse. Endocrinology. 2005;146(9):3686–92.

34. Hidalgo-Diaz C, Castano JP, Lopez-Pedrera R, Malagon MM, Garcia-Navarro S, Gracia-Navarro F. A modulatory role for substance P on the regulation of luteinizing hormone secretion by cultured porcine gonadotrophs. Biology of reproduction. 1998;58(3):678–85.

35. Hidalgo-Diaz C, Malagon MM, Garcia-Navarro S, Luque RM, Gonzalez de Aguilar JL, Gracia-Navarro F, et al. Role of Ca2+ in the secretory and biosynthetic response of porcine gonadotropes to substance P and gonadotropin-releasing hormone. Regul Pept. 2003;116(1-3):43–52.

36. Kerdelhue B, Tartar A, Lenoir V, el Abed A, Hublau P, Millar RP. Binding studies of substance P anterior pituitary binding sites: changes in substance P binding sites during the rat estrous cycle. Regul Pept. 1985;10(2-3):133–43.

37. Shamgochian MD, Leeman SE. Substance P stimulates luteinizing hormone secretion from anterior pituitary cells in culture. Endocrinology. 1992;131(2):871–5.

38. Kinsey-Jones JS, Grachev P, Li XF, Lin YS, Milligan SR, Lightman SL, et al. The inhibitory effects of neurokinin B on GnRH pulse generator frequency in the female rat. Endocrinology. 2012;153(1):307–15.

39. Leon S, Barroso A, Vazquez MJ, Garcia-Galiano D, Manfredi-Lozano M, Ruiz-Pino F, et al. Direct Actions of Kisspeptins on GnRH Neurons Permit Attainment of Fertility but are Insufficient to Fully Preserve Gonadotropic Axis Activity. Sci Rep. 2016;6:19206.

40. Martin C, Navarro VM, Simavli S, Vong L, Carroll RS, Lowell BB, et al. Leptin-responsive GABAergic neurons regulate fertility through pathways that result in reduced kisspeptinergic tone. The Journal of neuroscience: the official journal of the Society for Neuroscience. 2014;34(17):6047–56.

41. Steyn FJ, Wan Y, Clarkson J, Veldhuis JD, Herbison AE, Chen C. Development of a methodology for and assessment of pulsatile luteinizing hormone secretion in juvenile and adult male mice. Endocrinology. 2013;154(12):4939–45.

42. Garcia-Galiano D, Pineda R, Roa J, Ruiz-Pino F, Sanchez-Garrido MA, Castellano JM, et al. Differential modulation of gonadotropin responses to kisspeptin by aminoacidergic, peptidergic, and nitric oxide neurotransmission. American journal of physiology Endocrinology and metabolism. 2012;303(10):E1252–63.

43. Garcia-Galiano D, van Ingen Schenau D, Leon S, Krajnc-Franken MA, Manfredi-Lozano M, Romero-Ruiz A, et al. Kisspeptin signaling is indispensable for neurokinin B, but not glutamate, stimulation of gonadotropin secretion in mice. Endocrinology. 2012;153(1):316–28.

